# Cognitive deficits and increases in creatine precursors in a brain-specific knockout of the creatine transporter gene *Slc6a8*

**DOI:** 10.1101/196063

**Authors:** Kenea C. Udobi, Amanda N. Kokenge, Emily R. Hautman, Gabriela Ullo, Julie Coene, Michael T. Williams, Charles V. Vorhees, Aloïse Mabondzo, Matthew R Skelton

## Abstract

Creatine transporter (CrT; SLC6A8) deficiency (CTD) is an X-linked disorder characterized by severe cognitive deficits, impairments in language, and an absence of brain creatine (Cr). In a previous study, we generated floxed *Slc6a8 (Slc6a8^flox^)* mice to create ubiquitous *Slc6a8* knockout *(Slc6a8^-/y^)* mice. *Slc6a8^-/y^* mice lacked whole body Cr and exhibited cognitive deficits. While *Slc6a8^-/y^* mice have a similar biochemical phenotype to CTD patients, they also showed a reduction in size and reductions in swim speed that may have contributed to the observed deficits. To address this, we created brain-specific *Slc6a8* knockout (bKO) mice by crossing *Slc6a8^Flox^* mice with *Nestin-cre* mice. bKO mice had reduced cerebral Cr levels while maintaining normal Cr levels in peripheral tissue. Interestingly, brain concentrations of the Cr synthesis precursor guanidinoacetic acid were increased in bKO mice. bKO mice had longer latencies and path lengths in the Morris water maze, without reductions in swim speed. In accordance with data from *Slc6a8^-/y^* mice, bKO mice showed deficits in novel object recognition as well as contextual and cued fear conditioning. bKO mice were also hyperactive, in contrast with data from the *Slc6a8^-/y^*mice. The results demonstrate that the loss of cerebral Cr is responsible for the learning and memory deficits seen in ubiquitous *Slc6a8^-/y^* mice.

## Introduction

The absence of the high-energy phosphate buffer creatine (Cr) in the brain leads to intellectual disability (ID), language impairment, and epilepsy (Van De Kamp *et al.*, 2013). There are currently three known causes of cerebral Cr deficiency, of which Cr transporter (CrT; *SLC6A8*) deficiency (CTD) is the most prevalent. CTD has been estimated to be one of the leading causes of X-linked ID (XLID) in males (Almeida *et al.*, 2006, Clark *et al.*, 2006, Rosenberg *et al.*, 2004); however, little is known about the mechanisms underlying the cognitive deficits. To better understand the underlying patholophysiology of CTD, we generated mice with exons 2-4 of the murine *Slc6a8* gene flanked with LoxP sites (*Slc6a8^Flox^*). The primary goal in developing this model is to produce a high-fidelity model of CTD that recapitulates as much of the phenotype of human CTD patients as possible. In our first study, ubiquitous *Slc6a8* knockout (*Slc6a8^-/y^*) mice were derived from *Slc6a8^Flox^* mice (Skelton *et al.*, 2011). Consistent with human CTD, the *Slc6a8^-/y^*mice show an absence of brain Cr and cognitive deficits. *Slc6a8^-/y^* mice also had reduced body mass and a near absence of Cr in most tissues, including the muscle (Skelton *et al.*, 2011). The reduction of muscle Cr in *Slc6a8^-/y^* mice may be inconsistent with the limited data from human CTD showing normal muscle Cr levels (Degrauw *et al.*, 2003, Pyne-Geithman *et al.*, 2004). Further, the reduced size and loss of muscle Cr could influence the behavioral deficits observed in the *Slc6a8^-/y^* mice, since many of these tasks require physical movement. For example, *Slc6a8^-/y^* mice swam slower than *Slc6a8^+/y^* mice in the Morris water maze, which may have affected navigating to the hidden platform (Skelton *et al.*, 2011). This confound could reduce the translational value of this animal. By creating *Slc6a8^Flox^* mice, we are able to assess potential confounds using tissuespecific expression of Cre-recombinase.

Recently, an additional mouse model of CTD has been created by flanking exons 5-7 of the *Slc6a8* gene with LoxP sites (Baroncelli *et al.*, 2014). Similar to the *Slc6a8(2-4)^-/y^*mice of Skelton et al. (2011), ubiquitous knockout (*Slc6a8(5-7)^-ly^*) mice from this line showed reduced body weight and deficits in novel object recognition compared with *Slc6a8^+/y^* mice. Similar to the Slc6a8^flox/^-Camk2a mice, *Slc6a8(5-7)^-/y^* mice showed Morris water maze deficits during the last three days of testing (Baroncelli *et al.*, 2014). In a subsequent study *Slc6a8(5-7)^flox/y^* mice were crossed to *Nestin-cre* mice to create brain specific knockout mice. These mice show deficits in novel object recognition and Y-maze performance (Baroncelli *et al.*, 2016). While this supports the hypothesis that the effects are centrally mediated, the lack of MWM testing in these mice still leaves the possibility that the larger spatial learning deficit seen in *Slc6a8(2-4)^-/y^* mice are due to motor deficits-a confound that must be resolved in order to properly model this disorder.

Using the floxed *Slc6a8* (*Slc6a8^flox/y^*;FLOX) mouse line from the Skelton et al. (2011) study, Kurosawa et al. created a partial *Slc6a8* brain knockout mouse using a *Camk2a-cre* expressing mouse (Kurosawa *et al.*, 2012). The *Camk2a-cre* mouse expresses Cre recombinase in excitatory neurons in the forebrain (Tsien *et al.*, 1996). Similar to the *Slc6a8^-/y^* mice, the Slc6a8^flox/^-Camk2a mice showed spatial learning and novel object deficits (Kurosawa *et al.*, 2012). Interestingly, the spatial learning deficits were observed only when the last three days of testing were analyzed separately; in comparison *Slc6a8^-/y^* mice showed a robust deficit throughout testing. This would suggest that there could be some motor contribution to the deficit seen in *Slc6a8^-/y^* mice; however, the selection of the *Camk2a* mouse is somewhat problematic. This is due to *Camk2a-cre* expression being limited to excitatory neurons in the forebrain; allowing for the possiblity that some Cr-mediated function was spared. The *Nestin* promoter drives Cre recombinase expression in neuronal precursor cells, creating a more complete brain-specific knockout (Tronche *et al.*, 1999). The differences in these Creexpressing mice are highlighted by a recent study that showed that the offspring from floxed *Cc2d1a* mice crossed to *Nestin-cre* mice died at birth while the offspring from floxed mice crossed with *Camk2a-cre* mice were viable (Oaks *et al.*, 2017). The use of *Nestin-cre* mice” allows for a more complete understanding of the contributions of brain Cr to behavioral performance.

The purpose of this study was to determine if the cognitive deficits seen in the *Slc6a8^-/y^* mice were due to a lack of cerebral Cr by testing brain-specific *Slc6a8* knockout (bKO) mice. The behavioral tests used in this study have been shown to be disrupted in *Slc6a8^-/y^* mice (Hautman *et al.*, 2014, Skelton *et al.*, 2011) and represent three distinct types of learning with different neuroanatomical basis. The results of this study show that bKO mice have Morris water maze deficits similar to those seen in the ubiquitous *Slc6a8^-/y^* mice without reductions in swim speed. In addition, reductions in novel object recognition and conditioned fear memory were observed in bKO mice. This would suggest that the deficits observed in the *Slc6a8^-/y^* mice are central cognitive impairments analogous to those seen in CTD patients.

## Methods

### Generation of bKO mice

The bKO mice were generated and maintained on a C57BL/6J background in a pathogen-free facility in microisolators. Female *Slc6a8*^FLOX/+^ mice, generated and genotyped as described (Skelton *et al.*, 2011), were mated to mice expressing Cre recombinase driven by the *Nestin* promoter (B6.Cg-Tg (Nes-cre)^1Kln/J^; Jackson laboratory, Bar Harbor, ME) to create the mice used in this study. Testing was done during the light phase on *Slc6a8^+/y^::Nes-Cre^-^* (WT), bKO, *Slc6a8^+/y^::Nes-Cre^+^* (Nestin), and FLOX mice. Female mice were not used as heterozygous female *Slc6a8^+/−^* mice did not show reduced size or swim speed deficits (Hautman *et al.*, 2014). No more than one mouse per genotype was taken from a litter. All procedures on mice were performed at the CCRF vivarium which is fully accredited by AAALAC International and protocols were approved by the Institutional Animal Care and Use Committee. Lights were on a 14:10 light:dark cycle, room temperature was maintained at 19±1° C, and food (NIH-007) and filtered, sterilized water were available *ad libitum*. All institutional and national guidelines for the care and use of laboratory animals were followed.

### Behavioral Testing

Adult (between 60-90 days old) male mice of each genotype (n=14 WT; 15 bKO; 23 FLOX; 12 Nestin) were used. Mice were weighed on the first day of testing. Testing occurred in the order of presentation with 1 day between new tasks.

### Locomotor activity

Spontaneous locomotor activity (Brooks & Dunnett, 2009) was tested in automated activity chambers, Photobeam Activity System (PAS) – Open-Field (San Diego Instruments, San Diego, CA) for 1 h. Chambers were 40 cm (W) × 40 cm (D) × 38 cm (H) with 16 LED-photodetector beams in X and Y planes. Photocells were spaced 2.5 cm apart. Mice were tested for 1 h. The dependent measure was total number of photobeams interruptions.

### Morris water maze

The MWM is a test of spatial learning and reference memory (Vorhees & Williams, 2006); mice were tested as described (Skelton *et al.*, 2011). The tank was 122 cm in diameter, white, and filled with room temperature (21 ± 1°C) water. Prior to hidden platform testing, visible platform training (cued) was conducted for 6 days. During this phase, a 10 cm diameter platform with an orange ball mounted above the water on a brass rod was placed in one quadrant. On day 1, mice were given 6 trials with identical start and platform positions as training for the task. Mice were then given 2 trials per day for 5 days with the start and platform positions randomized and curtains closed around the pool to obscure distal cues to assess proximal cue learning.

Hidden platform trials were conducted in 2 phases; each phase consisted of 4 trials per day for 6 days followed by a single probe trial without the platform on day 7. Platform diameters were 10 cm during acquisition and 7 cm during reversal (platform located in the opposite quadrant). Reducing the size of the platform between acquisition and reversal increases the difficulty of the task and may be used to uncover spatial deficits (Vorhees & Williams, 2006). The trial limit was 90 s and the intertrial interval (ITI) was ~10 s. Mice that failed to find the platform were placed on the platform and given the same 10 s ITI. Performance was measured using ANY-maze^®^ software (Stoelting Company, Wood Dale, IL).

### Novel object recognition

Novel object recognition (NOR) is a test of incidental learning and memory (Clark *et al.*, 2000). Mice were tested in the ANY-box apparatus (Stoelting Company, Wood Dale, IL). First, mice were habituated to the arena (40×40 cm) for 2 days (10 min/day); next they were then habituated to two identical objects (10 min/day) for 2 days to reduce neophobia (Podhorna & Brown, 2002). On the fifth day, mice were presented with two new identical objects until 30 s of observation time between objects was accrued. One hour later, memory was tested by presenting the mouse with an identical copy of one of the familiar objects along with a novel object. Time exploring an object was defined as entry into a 2 cm zone around the object. Performance was measured using ANY-maze^®^ software. A discrimination index was calculated: time observing the novel object was subtracted from the time observing the familiar object, divided by total object observation time.

### Conditioned fear

Conditioned fear was assessed as described with modification (Peters *et al.*, 2010). On day 1, mice were placed in a chamber for 10 min before exposure to three tone-footshock pairings (82 dB, 2 kHz, 30 s on/off cycle). Each pairing consisted of the 30 s tone accompanied during the last second by a scrambled footshock (0.3 mA for 1 s) delivered through the floor. On the second day, mice were returned to the chamber with no tone or shock as a test of contextual fear. On the third day, mice were placed in the chamber with a novel floor. Following 3 min of acclimation, the tone was presented for 3 min. On day 2 and 3 freezing behavior was scored. Freezeframe software and Coulbourn test chambers were used (Coulbourn Instruments, Allentown, PA). Percent time freezing was analyzed.

### Creatine and guandinoacetic acid (GAA) determination

Following behavioral testing a subset of mice were sacrificed by decapitation. Brains were divided into hemispheres and flash frozen. Cr and GAA content were assayed in brain, heart, and kidney using a UPLC-MS/MS method(Trotier-Faurion *et al.*, 2013). First, 20 μL from each brain homogenate (diluted 1/10) was spiked with 5 μL of internal standard solution (creatine-D3-0.5 μg/mL) and 175μl of water were added to reach 200μl final volume. For liquid-liquid extraction, 400 μL ethanol and 400 μL hexane were added and the mixture was centrifuged at 4 °C for 15 min (10000g). Then, the upper phase was discarded and the hexane step was repeated one more time. Finally, the lower phase was transferred into a clean tube and evaporated to dryness under a gentle stream of nitrogen at 40°C. Then the samples were added with 200 μL of hydrogen chloride −1-butanol (≥99% HPLC Sigma-Aldrich, France) and placed in a Thermomixer at 65 °C for 15 min followed by an evaporation step. The residue was reconstituted with 100 μL of water injected into the LC-MS/MS system. The analytical method was performed with a binary pumps LC-30AD (Shimadzu Nexera X2) and compounds (creatine, GAA, creatine-D3) were separated by injecting extracted samples (10μL) on a C18 symmetry column-Waters (50 × 2.1 mm), with a 3.5 μm particles size maintained at 30 °C. The mobile phase consisted of phase A (0.1% formic acid in water) and phase B (0.1% formic acid in acetonitrile) on isocratic mode. Each analysis was carried out for 8 min and the flow rate of 0.4ml/min was used for the sample analysis. Detection was performed with a triple quadrupole mass spectrometer TSQ Quantum Ultra (Thermo) equipped with electrospray ionization source (ESI) in the positive ion mode. The equipment was set up in a multiple reaction monitoring (MRM) mode and the optimized setting parameters were: ion spray Voltage 3000V, source temperature 350 °C, sheath gas pressure 23, followed transitions for quantification: Creatine derivative: 188.18 → 90.20; 132.07 and 146.15 (Collision energy: 20); Creatine-D3 derivative: 190.93 →93.21; 134.87 and 149.23 (Collision energy: 20); GAA derivative: 174.09 → 72.75; 100.75 and 117.75 (Collision energy: 20). Results were recorded by Xcalibur and TSQ Tune Master (Thermo). The amount of creatine and GAA were standardized to the amount of protein in each homogenate tissue by the Bradford technique following the manufacturer’s instruction (Sigma-Aldrich, France).

### Statistics

Data were analyzed using one-way ANOVA except when there were repeated measurements on the same mouse (interval or day) in which case they were analyzed by repeated-measures ANOVA. Significant effects were analyzed using the method of Benjamini, Krieger, and Yekuteli (Benjamini *et al.*, 2006). Data were analyzed using GraphPad Prism.

## Results

### Body Weight

Mice were weighed on the first day of testing. No significant differences were observed in body weight between WT (26.4±0.6 g), FLOX (25.6±0.6 g), Nestin (25.3±1.0 g) and bKO (25.3±0.6 g) mice.

### Cr and GAA levels

#### Brain

There were main effects of gene for Cr (F(3,13)=39.94, p<0.001) and GAA (F(3,13)=4.3, p<0.05) levels (Figure 1A & B). Significant reductions in Cr were observed in bKO mice compared with WT, FLOX, and Nestin mice. There was a slight reduction in Cr levels in Nestin mice compared with WT mice (p<0.05) but not in FLOX mice. No differences were observed between FLOX and WT mice. GAA levels were increased in bKO mice compared with the control groups (p<0.05). No differences were observed between control groups for GAA.

**Figure 1.**
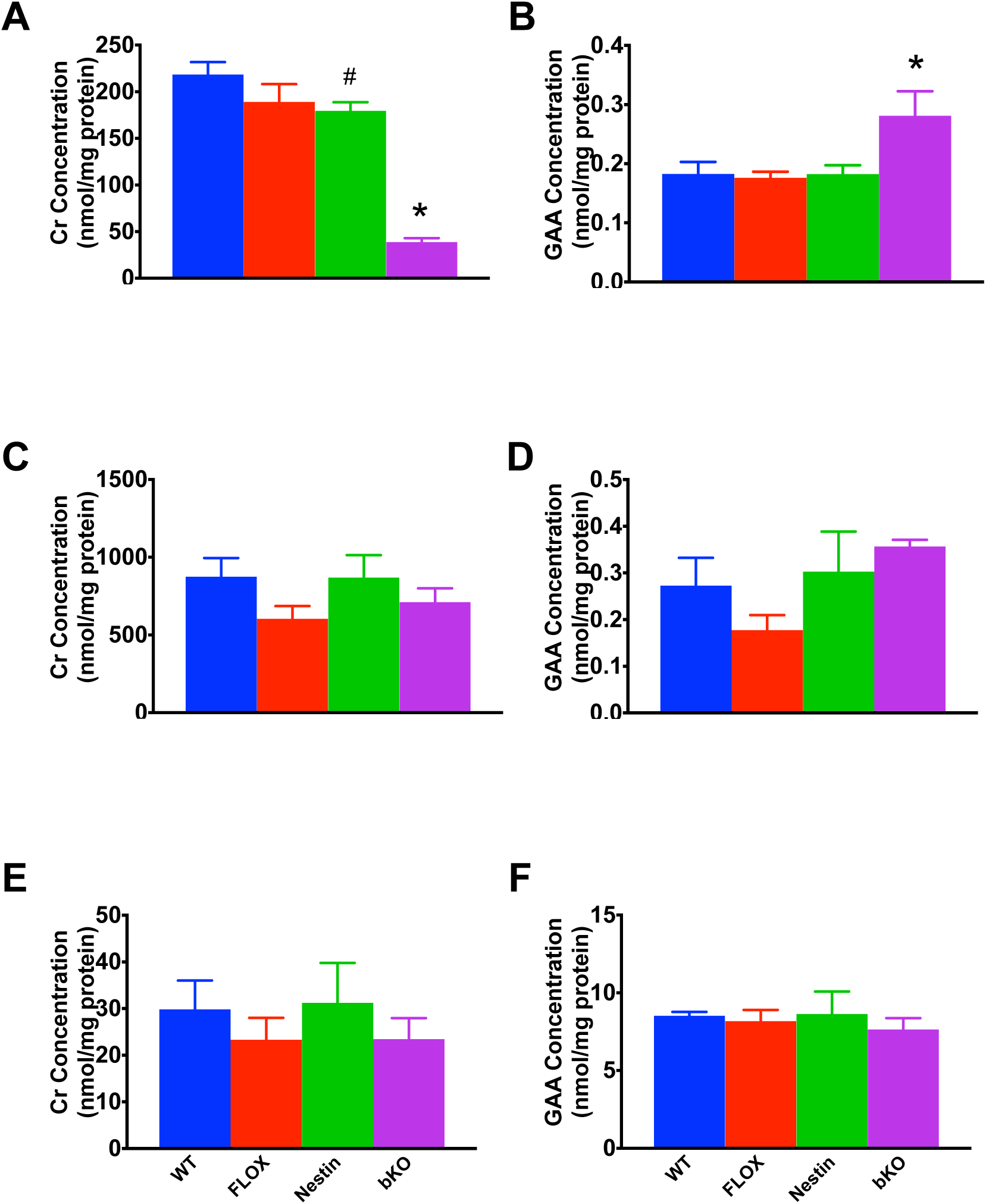
Creatine and GAA levels in the brain, heart and kidney. A) BKO mice show reduced brain Cr levels compared with all control groups. Nestin mice had lower Cr levels than WT mice. B) GAA levels are increased in the brain of bKO mice. In the heart and kidney, Cr (C and E respectively) and GAA (D and F) levels are unchanged. N=5/group Mean ±SEM *p<0.05 vs WT, FLOX, and NESTIN, #p<0.05 vs WT

#### Peripheral tissues

There was no effect of gene on Cr or GAA levels in the heart (Figure 1C & D). There was no effect of gene on Cr or GAA levels in the kidney (Figure 1E & F). There was no effect of gene on Cr or GAA levels in the muscle or liver (not shown for clarity).

### Locomotor Activity

There was a significant main effect of gene (F(3,54)=21.9; p<0.001; Figure 2) for total activity. A gene x time interaction (F(33, 594)=2.0, p<0.01) was observed. bKO mice showed increased activity compared with all other groups (p<0.05) at all times. FLOX mice showed hyperactivity compared with WT and Nestin mice at 15, 35, and 60 min. No differences were observed between WT and Nestin groups.

**Figure 2.**
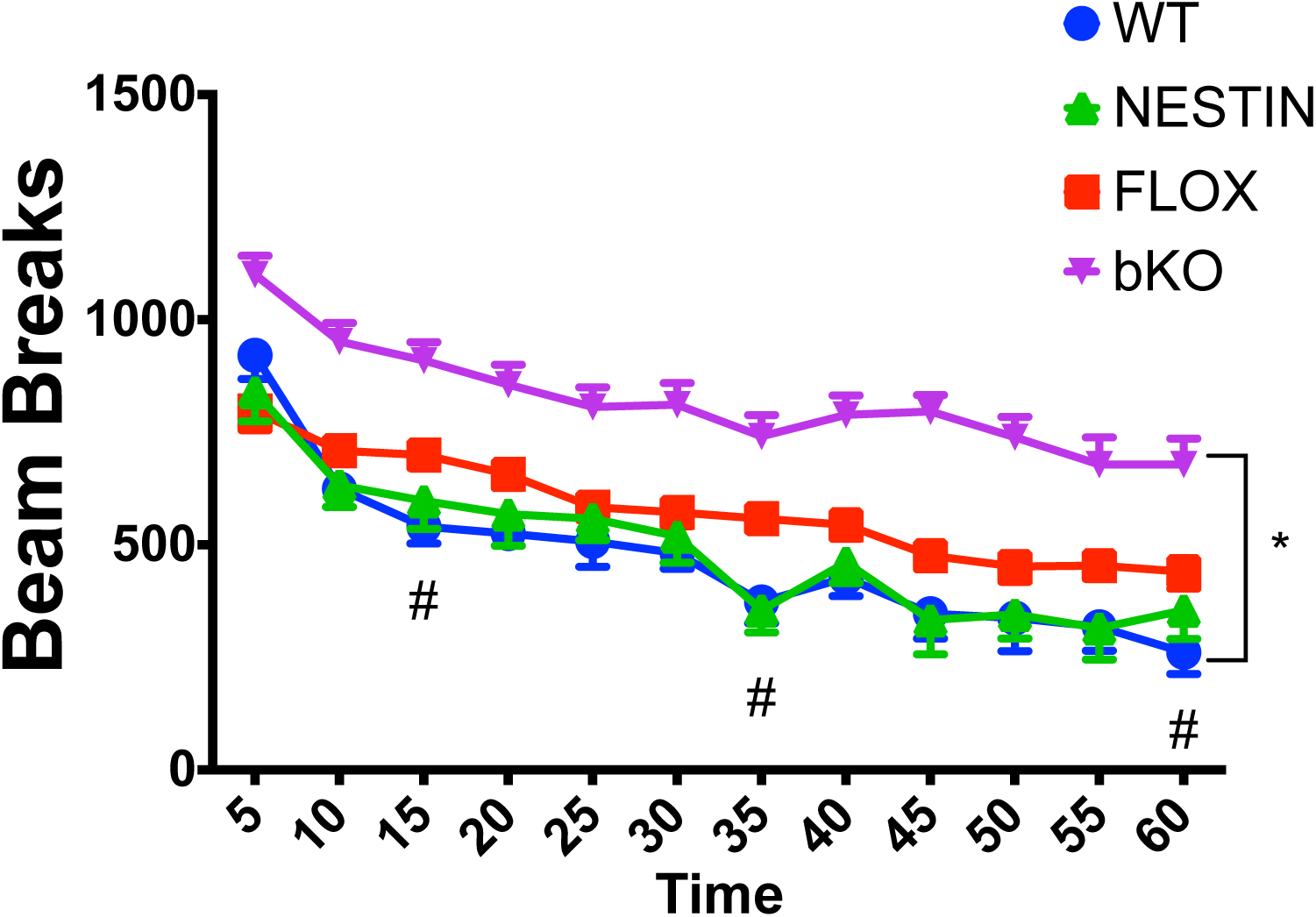
Hyperactivity in *Slc6a8* bKO mice. bKO mice show increased activity compared with FLOX and WT mice. FLOX mice showed increased activity compared with WT mice at 15 and 35 minutes. Data are presented as Mean ±SEM *p<0.05 vs FLOX, Nestin & WT, #p<0.05 vs WT & Nestin

### Morris water maze

On day-1 cued training, there was trend towards a main effect of gene (F(3,73)=2.6, p<0.07). An increase in latency was seen on days 2-6 (main effect of gene: F (3,73)=6.03, p<0.01). No differences were observed between the control groups.

For hidden platform testing, latency and path length were well correlated (r=0.832, p<0.001), therefore, path length is presented to remain consistent with our previous studies. During acquisition, bKO mice had an increased path length compared with control groups (F (3,72)=13.4, p<0.001; all post-hoc tests p<0.01 vs control groups; Figure 3A). No differences were observed between control groups. There were no significant effects of gene on swim speed. During the acquisition probe trial, bKO mice had a greater average distance from the former platform site compared with the WT mice (main effect of gene (F(3,72)=5.0, p<0.001; bKO vs WT p<0.05, Figure 3C). Learning curves for the WT vs bKO mice are shown in Figure 3E.

**Figure 3.**
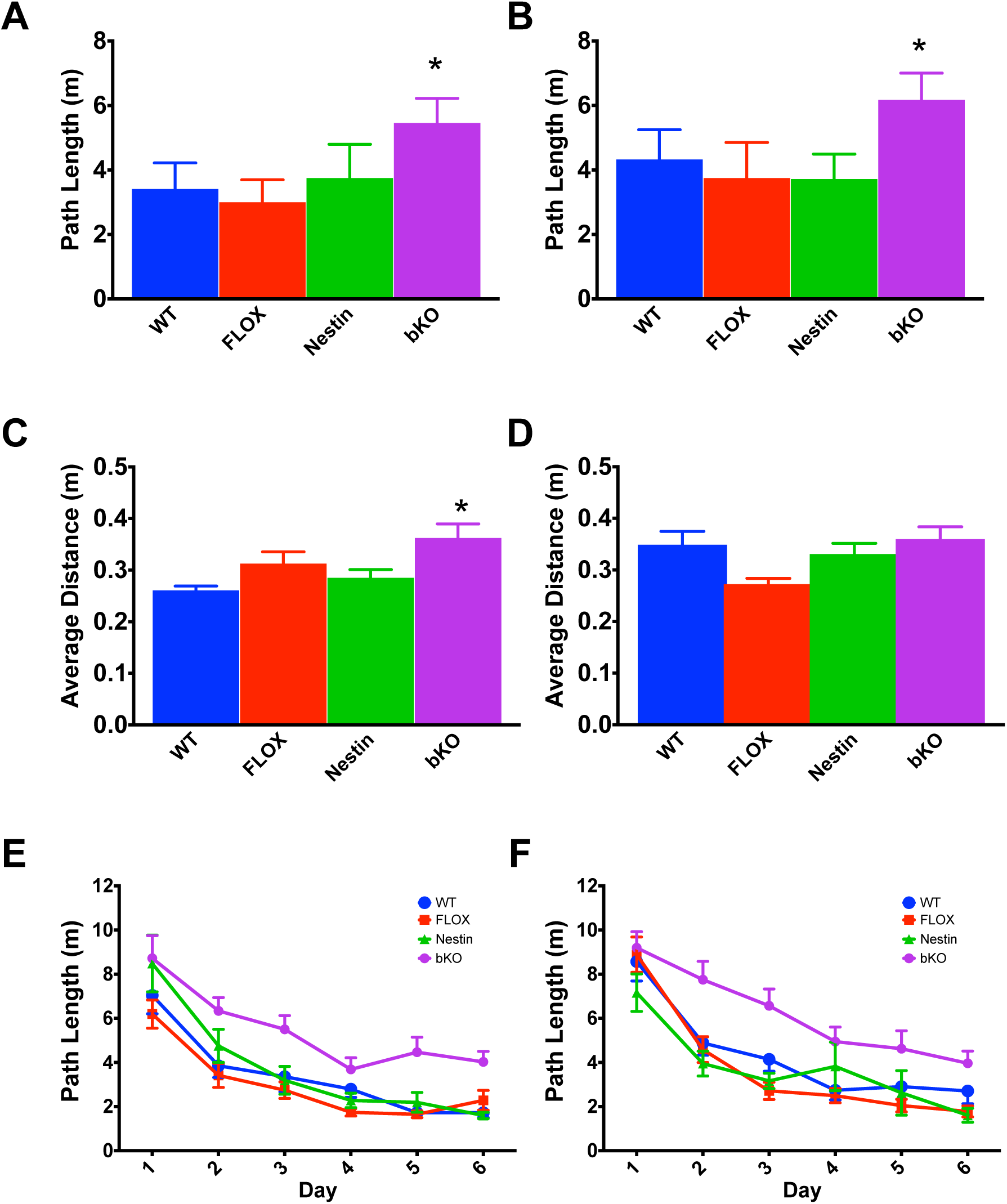
Spatial learning deficits in the MWM. bKO mice have increased path length to the platform during the acquisition (A) and reversal (B) phases of the MWM compared with WT and FLOX mice. On probe trials, bKO mice showed a higher average distance from the platform compared with controls during the acquisition phase (C) but not during the reversal phase (D). Learning curves by day for the bKO and WT mice during the acquisition (E) and reversal (F) hidden platform trials. Data are presented as Mean ±SEM **p<0.01 vs FLOX & WT; *p<0.05 vs FLOX & WT

During reversal, there was a main effect of gene for path length (F(3,69)=10.2, p<0.001; Figure 3B). bKO mice travelled a longer path to the platform compared with the control groups (p<0.05). No differences were observed between control groups. There were no differences in swim speed during reversal. During the reversal probe trial, there was a main effect of gene (F(3,69)=4.3, p<0.01; Figure 3D). FLOX mice had shorter average distance to the target site compared with WT and bKO mice (p<0.05). No other differences were observed. Learning curves for WT vs bKO mice are shown in Figure 3F.

### Novel Object Recognition

For time spent observing the novel object, there was a main effect of gene (F(3,56)=7.7, p<0.001; Figure 4). Post-hoc tests showed that bKO mice spent less time with the novel object compared with control groups (p<0.05); the control groups did not differ from each other.

**Figure 4.**
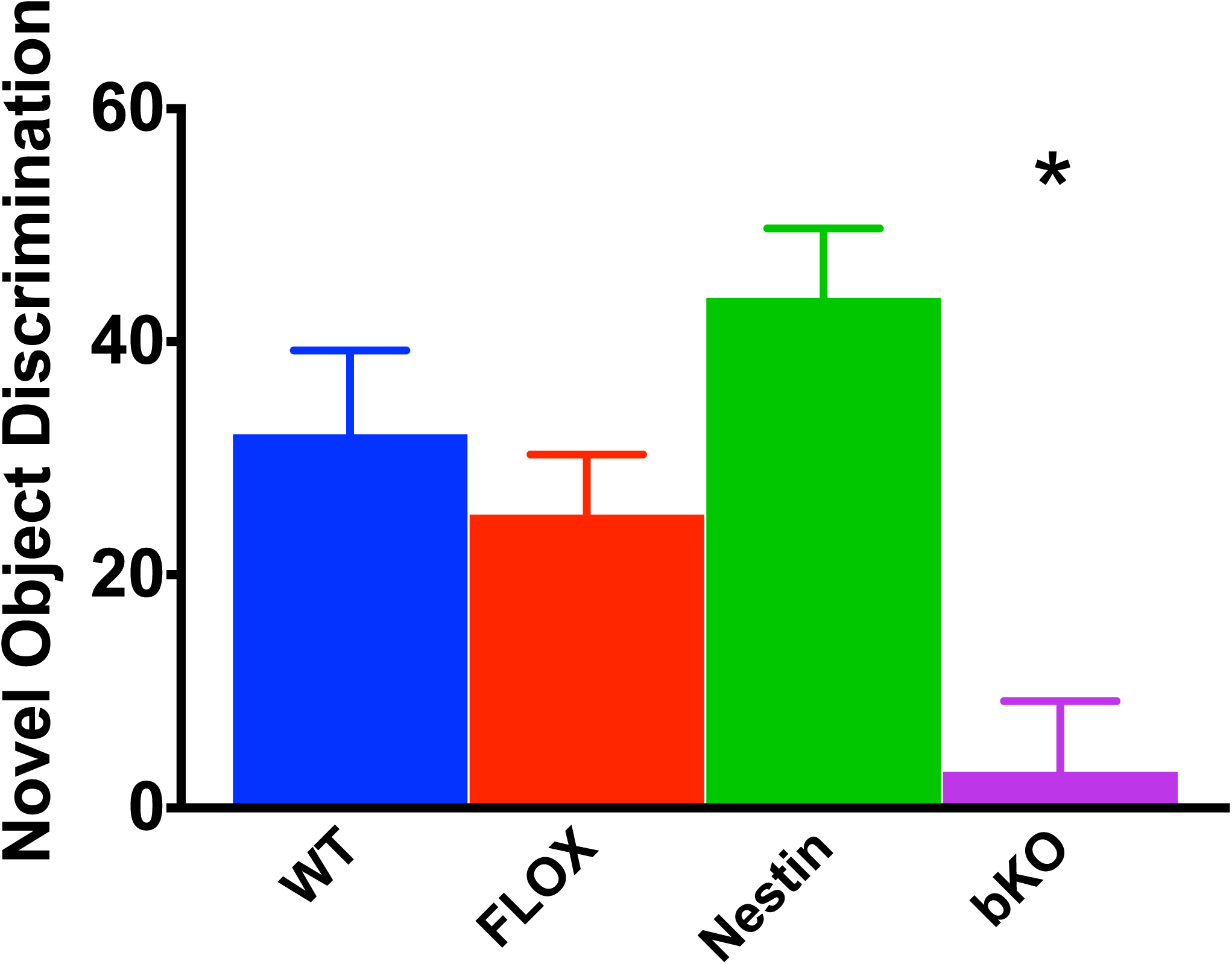
Object recognition memory deficits. bKO mice had a reduced discrimination index for the novel object compared with the control groups. Data are presented as Mean ±SEM *p<0.05 vs WT, FLOX & Nestin

### Conditioned Fear

On day 1, mice were conditioned to the tone-footshock pairing. On day 2, there was a main effect of gene (F (3, 60) =7.7, p<0.01, Figure 4) that was the result of bKO mice spending less time freezing than WT and FLOX mice in the training context (contextual fear). Nestin mice did not show a significant difference from other groups. On day 3, there was a main effect of gene (F(3,64) = 10.3, p<0.001,Figure 5);bKO mice spent less time freezing to the tone (cued fear) than WT and FLOX mice.

**Figure 5.**
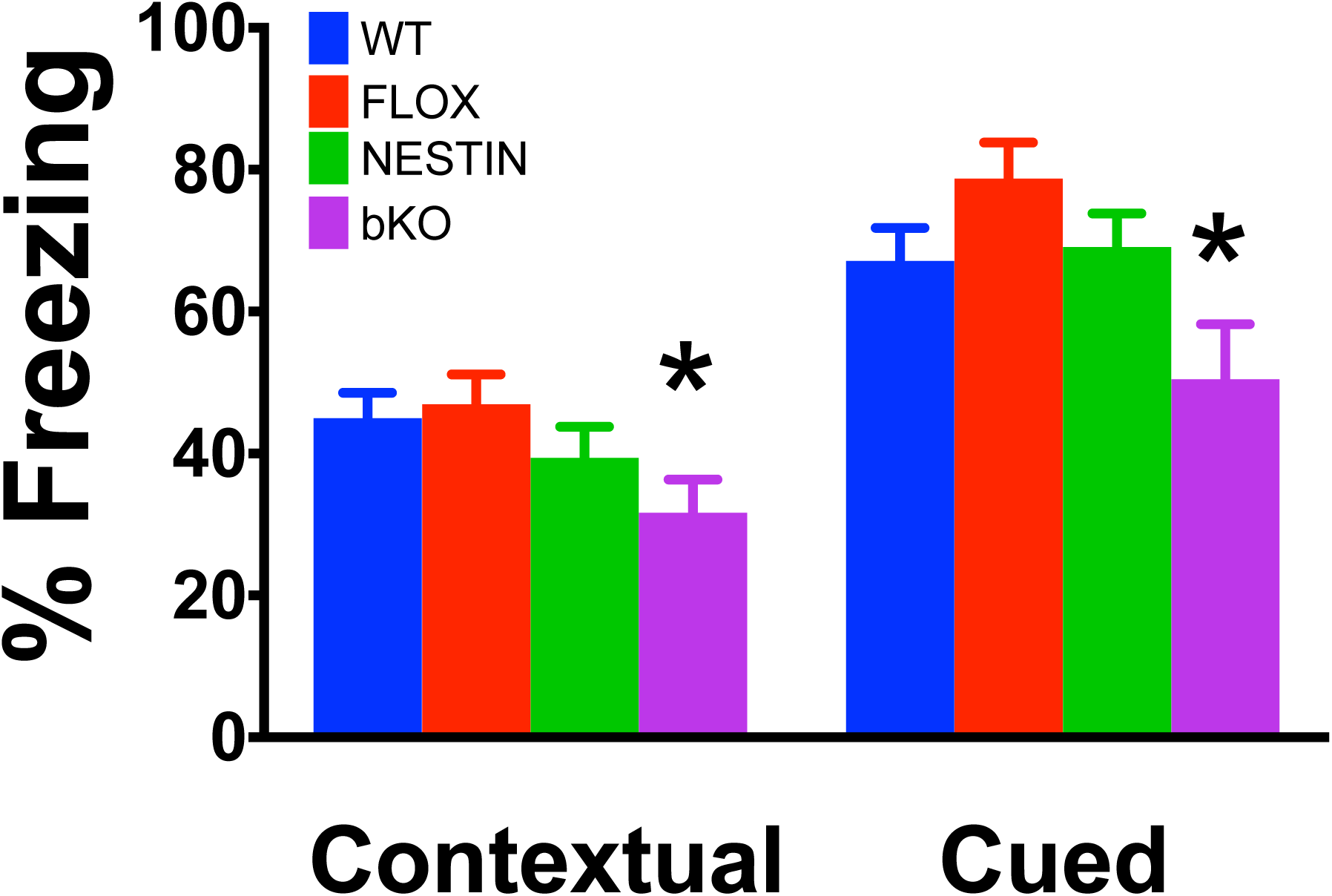
Contextual and conditioned fear deficits in *Slc6a8* bKO mice. bKO mice showed reduced freezing compared with control groups during both the contextual and cued phases of testing. Data are presented as Mean ±SEM **p<0.01 vs FLOX & WT.

## Discussion

The study was conducted to determine if cognitive deficits observed in ubiquitous *Slc6a8^-/y^* mice were influenced by sensorimotor deficits caused by the lack of peripheral Cr. The results show that *Nestin-cre::Slc6a8^FLOX/y^* mice have robust learning and memory deficits, similar to *Slc6a8^-/y^* mice but in the absence of body weight reductions or slow swimming. This would suggest that the effects seen in *Slc6a8^-/y^* mice are due to a lack of cerebral Cr.

Similar to the ubiquitous *Slc6a8^-/y^* mice, bKO mice show learning and memory deficits in the MWM, NOR, and conditioned fear, suggesting that these mice have wide-ranging cognitive deficits. This supports the hypothesis that these mice are a high-fidelity model since CTD patients appear to show a global cognitive delay (Van De Kamp *et al.*, 2013). In contrast to *Slc6a8^-/y^* mice, the MWM deficits appeared without differences in swim speed. Interestingly, the bKO mice had deficits in the cued platform phase, which is a control procedure for sensorimotor deficits. However, given a lack of swim speed changes during hidden platform testing, it is unlikely that the cued effect is carrying over to the spatial learning phases of the test. It is also unlikely that the hyperactivity seen in the bKO mice played a role in MWM performance, as land based hyperactivity does not often translate to increases in exploration in water-based tasks (Vorhees & Williams, 2006). In addition, female *Slc6a8^+/−^* mice were hyperactive and did not show deficits in NOR or conditioned fear. Further, the data from all models used to date have provided converging evidence of cognitive deficits in both the presence and absence of hyperactivity. Taken together, it is possible but unlikely that the hyperactivity plays a significant role in the cognitive deficits observed.

Attention deficit/hyperactivity disorder has been reported in many CTD patients (Dunbar *et al.*, 2014). Ubiquitous *Slc6a8^-/y^* mice showed initial hypoactivity followed by no difference in an open-field test compared with WT mice. In contrast, bKO mice were hyperactive. In these studies, a photobeam system was used to determine movement. It is possible that the smaller stature of the *Slc6a8^-/y^*mice lead to fewer beam interruptions but the exact cause is unknown. The presence of hyperactivity in these mice represent a new finding in our mice that is present in the CTD population, making the bKO mice a viable model for certain aspects of CTD.

*Slc6a8^-/y^* mice are smaller than *Slc6a8^+/y^* mice which is similar to mice that are deficient in Cr synthesis (Choe *et al.*, 2013, Schmidt *et al.*, 2004). The reduction in size is in agreement with human data, as recent case reports show that hypotonia is a phenotype in many CTD patients (Dunbar *et al.*, 2014), suggesting that the reduced size of the *Slc6a8^-/y^* mouse matches effects found in many cases of CTD. Interestingly, *Slc6a8^-/y^* mice have increased body fat percentages compared with *Slc6a8^+/y^* mice (Perna *et al.*, 2016), which could suggest a loss of muscle mass in the mice. Both CNS and peripheral systems contribute to body mass. The finding of normal body mass in the bKO mice suggests that body fat changes are peripherally-mediated and not due to a loss of hypothalamic Cr. This is further supported by a recent study where adipose-tissue specific deletion of the Cr-synthesis precursor *Arginine:glycine amidinotransferase* (*Agat* or *Gatm*) increased body fat in mice (Kazak *et al.*, 2017).

Brain Cr levels were reduced in the bKO mice while Cr levels in other tissues were spared. This suggests that there was no carry-over effect using the *Nestin-cre* in other tissues. There was a small decrease in brain Cr in the *Nestin-cre* mice. The reduction in brain Cr did not appear to influence the behavior of the animals. This is not surprising since female *Slc6a8^+/−^*mice only have mild cognitive impairments despite a 50% reduction in brain Cr levels (Hautman *et al.*, 2014). Interestingly, there was an increase in the Cr synthesis precursor GAA in the brains of the bKO mice. This supports the hypothesis that the rodent brain can synthesize Cr and uses the *Slc6a8* to shuttle Cr intermediates between cells (Braissant *et al.*, 2011). With a few exceptions, elevations in GAA levels are not observed in the urine and CSF of CTD patients (Van De Kamp *et al.*, 2013, Van De Kamp *et al.*, 2012a, Van De Kamp *et al.*, 2012b), This does not exclude the possibility of increased GAA in the brain of CTD patients, but these findings do warrant further study into determine if and how the brain synthesizes Cr and what impact this has for CTD patients. CTD patients supplemented with Cr-synthesis precursors arginine and glycine did not show an increase in cerebral Cr levels (Jaggumantri *et al.*, 2015, Valayannopoulos *et al.*, 2012). If this system can be better understood, it is possible that future treatment could involve enhancing this treatment protocol to somehow initiate endogenous Cr synthesis in the brain.

Choosing the proper model for understanding CTD is of utmost importance for understanding this disorder and developing treatments. The bKO mice used in this study represent the fifth genetic model of CTD examined. Three have come from the *Slc6a8^FLOX/y^* mice used in this study and two from the *Slc6a8(5-7)^FLOX/y^* mice. In addition to the bKO mice from this study and the *Slc6a8^-/y^* mice from Skelton et al (2011), Kurosawa et al (2012) generated *Camk2a-cre:: Slc6a8^FLOX/y^* mice, creating a forebrain-specific *Slc6a8* knockout (Tsien *et al.*, 1996). These mice showed a reduction in object recognition memory but modest deficits in the MWM that were only seen on when the last three days of testing were analyzed separately from the overall ANOVA. In contrast, the learning curves of the WT and bKO mice (Figure 3E) show that the deficits began on day 1 in the acquisition phase. This is similar to what is seen in the *Slc6a8^-/y^*mice, suggesting that the bKO mice recapitulates the learning deficits seen in the ubiquitous *Slc6a8^-/y^* mouse and in CTD. Along with the selective elimination of the *Slc6a8* gene, the smaller spatial learning deficit seen in the Kurosawa et al. paper may be due to the presence of some Cr during brain development in the *Camk2a-cre*:: *Slc6a8^FLOX/y^* mice. *Camk2a-cre* expression is not observed until approximately postnatal day 3 (Burgin *et al.*, 1990, Guo *et al.*, 2000), allowing for the presence of Cr during critical periods of brain development. Further, since the elimination of Cr is ~3% per day (Wyss & Kaddurah-Daouk, 2000), Cr could be present at high levels for several weeks even after *Slc6a8* deletion in the *Camk2a-cre:: Slc6a8^FLOX/y^* mice. The importance of Cr in brain development has been highlighted in a recent study showing that the loss of Cr kinase leads to aberrant neuronal development (Fukumitsu *et al.*, 2015). In addition, maternal Cr supplementation in rats enhanced hippocampal neuron development and improved long-term potentiation (Sartini *et al.*, 2016). Taken together, these data suggest that using *Nestin-cre* mice to generate brain-specific *Slc6a8* disruption is more comparable to the ubiquitous *Slc6a8^-/y^* mice.

Baroncelli et al. (2016) showed that a *Nestin-cre* generated brain-specific knockout of exons 5-7 of the *Slc6a8* gene resulted in novel object recognition and spontaneous alternation deficits. Ubiquitous deletion of the same region results in Morris water maze deficits, but only when the last 3 days of testing were examined separately (Baroncelli *et al.*, 2014). Interestingly, this deficit did not fully replicate in a subsequent study when similarly-aged mice showed a deficit only on day 5 of testing (Baroncelli *et al.*, 2016). This would suggest that both our ubiquitous and our bKO mice have more severe spatial learning deficits than the ubiquitous 5-7 deletion. The brain-specific knockout of exons 5-7 mice were not evaluated in the Morris water maze. The difference in spatial learning between the two *Slc6a8* knockouts could also be explained by differences in the test apparatus or procedures; for example, platform to arena size ratio in these studies differ. We used a 144:1 tank to platform ratio for the acquisition phase (Hautman *et al.*, 2014, Skelton *et al.*, 2011), while the Baroncelli studies had a 93.4:1 tank to platform ratio. Smaller search ratios simplify the task (Vorhees & Williams, 2006), perhaps making the lower ratio apparatus less sensitive. Rectifying these differences could provide additional information about this disorder by relating different mutation to known variants in human CTD. This could prove to be difficult as there is a great deal of heterogeneity in CTD-causing mutations (Van De Kamp *et al.*, 2013).

In conclusion, the results of this study show that the cognitive deficits in ubiquitous *Slc6a8^-/y^* mice are attributable to changes in cognitive function and are not caused by being smaller or having motor deficits. Based on this, we suggest that the *Slc6a8^-/y^* mouse is the best model of human CTD. While smaller stature is not always observed in human CTD, all CTD patients have ubiquitous deletion of the gene. Brain-specific models are generated using neuronal precursors that likely allow for the expression of the *Slc6a8* gene at the blood brain barrier (BBB). At the BBB, *Slc6a8* is expressed in microcapillary endothelial cells and not on astrocytic feet that encompass the majority of the BBB (Braissant & Henry, 2008). These cells derive from a mesodermal lineage and any neuronal model of Cre expression would not target them, allowing for *Slc6a8* expression at the BBB. Taken together, the bKO data support that, regardless of the model used, ubiquitous *Slc6a8* knockout mice are the preferred model of CTD; whereas the bKO models should be limited to studies to provide evidence that the ubiquitous model is not confounded. Therefore, the ubiquitous model should be utilized to develop potential therapeutics for this disorder.

## References

Almeida, L.S., Rosenberg, E.H., Verhoeven, N.M., Jakobs, C. & Salomons, G.S. (2006) Are cerebral creatine deficiency syndromes on the radar screen? Future Neurol., 1, 637–649.

Baroncelli, L., Alessandri, M.G., Tola, J., Putignano, E., Migliore, M., Amendola, E., Gross, C., Leuzzi, V., Cioni, G. & Pizzorusso, T. (2014) A novel mouse model of creatine transporter deficiency. F1000Res, 3, 228.

Baroncelli, L., Molinaro, A., Cacciante, F., Alessandri, M.G., Napoli, D., Putignano, E., Tola, J., Leuzzi, V., Cioni, G. & Pizzorusso, T. (2016) A mouse model for creatine transporter deficiency reveals early onset cognitive impairment and neuropathology associated with brain aging. Hum Mol Genet.

Benjamini, Y., Krieger, A.M. & Yekutieli, D. (2006) Adaptive linear step-up procedures that control the false discovery rate. Biometrika, 93, 491–507.

Braissant, O. & Henry, H. (2008) AGAT, GAMT and SLC6A8 distribution in the central nervous system, in relation to creatine deficiency syndromes: A review. J.Inherit.Metab Dis.

Braissant, O., Henry, H., Beard, E. & Uldry, J. (2011) Creatine deficiency syndromes and the importance of creatine synthesis in the brain. Amino.Acids.

Brooks, S.P. & Dunnett, S.B. (2009) Tests to assess motor phenotype in mice: a user’s guide. Nat.Rev.Neurosci., 10, 519–529.

Burgin, K.E., Waxham, M.N., Rickling, S., Westgate, S.A., Mobley, W.C. & Kelly, P.T. (1990) In situ hybridization histochemistry of Ca2+/calmodulin-dependent protein kinase in developing rat brain. J Neurosci, 10, 1788–1798.

Choe, C.U., Nabuurs, C., Stockebrand, M.C., Neu, A., Nunes, P., Morellini, F., Sauter, K., Schillemeit, S., Hermans-Borgmeyer, I., Marescau, B., Heerschap, A. & Isbrandt, D. (2013) Larginine:glycine amidinotransferase deficiency protects from metabolic syndrome. Hum Mol Genet, 22, 110–123.

Clark, A.J., Rosenberg, E.H., Almeida, L.S., Wood, T.C., Jakobs, C., Stevenson, R.E., Schwartz, C.E. & Salomons, G.S. (2006) X-linked creatine transporter (SLC6A8) mutations in about 1% of males with mental retardation of unknown etiology. Hum.Genet., 119, 604–610.

Clark, R.E., Zola, S.M. & Squire, L.R. (2000) Impaired recognition memory in rats after damage to the hippocampus. J Neurosci, 20, 8853–8860.

deGrauw, T.J., Cecil, K.M., Byars, A.W., Salomons, G.S., Ball, W.S. & Jakobs, C. (2003) The clinical syndrome of creatine transporter deficiency. Mol Cell Biochem, 244, 45–48.

Dunbar, M., Jaggumantri, S., Sargent, M., Stockler-Ipsiroglu, S. & van Karnebeek, C.D. (2014) Treatment of X-linked creatine transporter (SLC6A8) deficiency: systematic review of the literature and three new cases. Mol Genet Metab, 112, 259–274.

Fukumitsu, K., Fujishima, K., Yoshimura, A., Wu, Y.K., Heuser, J. & Kengaku, M. (2015) Synergistic action of dendritic mitochondria and creatine kinase maintains ATP homeostasis and actin dynamics in growing neuronal dendrites. J Neurosci, 35, 5707–5723.

Guo, H., Mao, C., Jin, X.L., Wang, H., Tu, Y.T., Avasthi, P.P. & Li, Y. (2000) Cre-mediated cerebellum- and hippocampus-restricted gene mutation in mouse brain. Biochem Biophys Res Commun, 269, 149–154.

Hautman, E.R., Kokenge, A.N., Udobi, K.C., Williams, M.T., Vorhees, C.V. & Skelton, M.R. (2014) Female mice heterozygous for creatine transporter deficiency show moderate cognitive deficits. J Inherit Metab Dis, 37, 63–68.

Jaggumantri, S., Dunbar, M., Edgar, V., Mignone, C., Newlove, T., Elango, R., Collet, J.P., Sargent, M., Stockler-Ipsiroglu, S. & van Karnebeek, C.D. (2015) Treatment of Creatine Transporter (SLC6A8) Deficiency With Oral S-Adenosyl Methionine as Adjunct to L-arginine, Glycine, and Creatine Supplements. Pediatr Neurol, 53, 360–363 e362.

Kazak, L., Chouchani, E.T., Lu, G.Z., Jedrychowski, M.P., Bare, C.J., Mina, A.I., Kumari, M., Zhang, S., Vuckovic, I., Laznik-Bogoslavski, D., Dzeja, P., Banks, A.S., Rosen, E.D. & Spiegelman, B.M. (2017) Genetic Depletion of Adipocyte Creatine Metabolism Inhibits Diet-Induced Thermogenesis and Drives Obesity. Cell Metab.

Kurosawa, Y., Degrauw, T.J., Lindquist, D.M., Blanco, V.M., Pyne-Geithman, G.J., Daikoku, T., Chambers, J.B., Benoit, S.C. & Clark, J.F. (2012) Cyclocreatine treatment improves cognition in mice with creatine transporter deficiency. J Clin Invest.

Oaks, A.W., Zamarbide, M., Tambunan, D.E., Santini, E., Di Costanzo, S., Pond, H.L., Johnson, M.W., Lin, J., Gonzalez, D.M., Boehler, J.F., Wu, G.K., Klann, E., Walsh, C.A. & Manzini, M.C. (2017) Cc2d1a Loss of Function Disrupts Functional and Morphological Development in Forebrain Neurons Leading to Cognitive and Social Deficits. Cereb Cortex, 27, 1670–1685.

Perna, M.K., Kokenge, A.N., Miles, K.N., Udobi, K.C., Clark, J.F., Pyne-Geithman, G.J., Khuchua, Z. & Skelton, M.R. (2016) Creatine transporter deficiency leads to increased whole body and cellular metabolism. Amino Acids, 48, 2057–2065.

Peters, J., Dieppa-Perea, L.M., Melendez, L.M. & Quirk, G.J. (2010) Induction of fear extinction with hippocampal-infralimbic BDNF. Science, 328, 1288–1290.

Podhorna, J. & Brown, R.E. (2002) Strain differences in activity and emotionality do not account for differences in learning and memory performance between C57BL/6 and DBA/2 mice. Genes Brain Behav, 1, 96–110.

Pyne-Geithman, G.J., DeGrauw, T.J., Cecil, K.M., Chuck, G., Lyons, M.A., Ishida, Y. & Clark, J.F. (2004) Presence of normal creatine in the muscle of a patient with a mutation in the creatine transporter: a case study. Mol.Cell Biochem., 262, 35–39.

Rosenberg, E.H., Almeida, L.S., Kleefstra, T., deGrauw, R.S., Yntema, H.G., Bahi, N., Moraine, C., Ropers, H.H., Fryns, J.P., DeGrauw, T.J., Jakobs, C. & Salomons, G.S. (2004) High prevalence of SLC6A8 deficiency in X-linked mental retardation. Am.J.Hum.Genet., 75, 97–105.

Sartini, S., Lattanzi, D., Ambrogini, P., Di Palma, M., Galati, C., Savelli, D., Polidori, E., Calcabrini, C., Rocchi, M.B., Sestili, P. & Cuppini, R. (2016) Maternal creatine supplementation affects the morpho-functional development of hippocampal neurons in rat offspring. Neuroscience, 312, 120–129.

Schmidt, A., Marescau, B., Boehm, E.A., Renema, W.K., Peco, R., Das, A., Steinfeld, R., Chan, S., Wallis, J., Davidoff, M., Ullrich, K., Waldschutz, R., Heerschap, A., De Deyn, P.P., Neubauer, S. & Isbrandt, D. (2004) Severely altered guanidino compound levels, disturbed body weight homeostasis and impaired fertility in a mouse model of guanidinoacetate N-methyltransferase (GAMT) deficiency. Hum Mol Genet, 13, 905–921.

Skelton, M.R., Schaefer, T.L., Graham, D.L., Degrauw, T.J., Clark, J.F., Williams, M.T. & Vorhees, C.V. (2011) Creatine transporter (CrT; Slc6a8) knockout mice as a model of human CrT deficiency. PLoS One, 6, e16187.

Tronche, F., Kellendonk, C., Kretz, O., Gass, P., Anlag, K., Orban, P.C., Bock, R., Klein, R. & Schutz, G. (1999) Disruption of the glucocorticoid receptor gene in the nervous system results in reduced anxiety. Nat Genet, 23, 99–103.

Trotier-Faurion, A., Dezard, S., Taran, F., Valayannopoulos, V., de Lonlay, P. & Mabondzo, A. (2013) Synthesis and biological evaluation of new creatine fatty esters revealed dodecyl creatine ester as a promising drug candidate for the treatment of the creatine transporter deficiency. J Med Chem, 56, 5173–5181.

Tsien, J.Z., Chen, D.F., Gerber, D., Tom, C., Mercer, E.H., Anderson, D.J., Mayford, M., Kandel, E.R. & Tonegawa, S. (1996) Subregion- and cell type-restricted gene knockout in mouse brain. Cell, 87, 1317–1326.

Valayannopoulos, V., Boddaert, N., Chabli, A., Barbier, V., Desguerre, I., Philippe, A., Afenjar, A., Mazzuca, M., Cheillan, D., Munnich, A., de Keyzer, Y., Jakobs, C., Salomons, G.S. & de Lonlay, P. (2012) Treatment by oral creatine, L-arginine and L-glycine in six severely affected patients with creatine transporter defect. J Inherit Metab Dis, 35, 151–157.

van de Kamp, J.M., Betsalel, O.T., Mercimek-Mahmutoglu, S., Abulhoul, L., Grunewald, S., Anselm, I., Azzouz, H., Bratkovic, D., de Brouwer, A., Hamel, B., Kleefstra, T., Yntema, H., Campistol, J., Vilaseca, M.A., Cheillan, D., D’Hooghe, M., Diogo, L., Garcia, P., Valongo, C., Fonseca, M., Frints, S., Wilcken, B., von der Haar, S., Meijers-Heijboer, H.E., Hofstede, F., Johnson, D., Kant, S.G., Lion-Francois, L., Pitelet, G., Longo, N., Maat-Kievit, J.A., Monteiro, J.P., Munnich, A., Muntau, A.C., Nassogne, M.C., Osaka, H., Ounap, K., Pinard, J.M., Quijano-Roy, S., Poggenburg, I., Poplawski, N., Abdul-Rahman, O., Ribes, A., Arias, A., Yaplito-Lee, J., Schulze, A., Schwartz, C.E., Schwenger, S., Soares, G., Sznajer, Y., Valayannopoulos, V., Van Esch, H., Waltz, S., Wamelink, M.M., Pouwels, P.J., Errami, A., van der Knaap, M.S., Jakobs, C., Mancini, G.M. & Salomons, G.S. (2013) Phenotype and genotype in 101 males with X-linked creatine transporter deficiency. J Med Genet.

van de Kamp, J.M., Jakobs, C., Gibson, K.M. & Salomons, G.S. (2012a) New insights into creatine transporter deficiency: the importance of recycling creatine in the brain. J Inherit Metab Dis.

van de Kamp, J.M., Pouwels, P.J., Aarsen, F.K., ten Hoopen, L.W., Knol, D.L., de Klerk, J.B., de Coo, I.F., Huijmans, J.G., Jakobs, C., van der Knaap, M.S., Salomons, G.S. & Mancini, G.M. (2012b) Long-term follow-up and treatment in nine boys with X-linked creatine transporter defect. J Inherit Metab Dis, 35, 141–149.

Vorhees, C.V. & Williams, M.T. (2006) Morris water maze: procedures for assessing spatial and related forms of learning and memory. Nature Protocols, 1, 848–858.

Wyss, M. & Kaddurah-Daouk, R. (2000) Creatine and creatinine metabolism. Physiol Rev., 80, 1107–1213.

